# Delay of punishment highlights differential vulnerability to developing addiction-like behavior toward sweet food

**DOI:** 10.1101/2023.10.09.559890

**Authors:** Marcello Solinas, Virginie Lardeux, Pierre-Marie LeBlanc, Jean-Emmanuel Longueville, Nathalie Thiriet, Youna Vandaele, Leigh Panlilio, Nematollah Jaafari

## Abstract

Resistance to punishment is commonly used as a measure of compulsive behavior in addiction-related processes. We recently developed a progressive shock strength (PSS) procedure in which individual rats can titrate the amount of punishment that they are willing to tolerate to obtain food rewards. Here, we investigated the effects of a range of delays (0-12 sec) on resistance to punishment measured by PSS break points. As expected from delay discounting principles, we found that delayed shock was less effective as a punisher, as revealed by higher PSS breakpoints. However, this discounting effect was not equally distributed in the population of rats, and the introduction of a delay highlighted the existence of two populations: rats that were sensitive to immediate punishment were also weakly sensitive to delay, whereas rats that were resistant to immediate punishment showed strong temporal discounting of delayed punishment. Importantly, shock-sensitive rats suppressed responding even in non-punishment sessions, and they differed from shock-resistant rats in anxiety-like behavior but not in sensitivity to pain. These results show that manipulation of temporal contingencies of punishment in the PSS procedure provides a valuable tool to identify individuals with a double vulnerability to addiction: low sensitivity to aversion and excessive discounting of negative future consequences. Conversely, the shock-sensitive population may provide a model of humans who are vulnerable to opportunity loss due to excessive anxiety.

---

Drug addiction is characterized by excessive motivation to consume the drug and continued use in spite of negative consequences [1]. The motivational aspects of addiction and the role of the reward system have been extensively studied [2–8]; however, the investigation of the behavioral and neurobiological mechanisms underlying the inability to stop drug use in face of adverse consequences has attracted the attention of the field only in the last decade [9–13]. Whereas these processes are intertwined, behavioral and neurobiological evidence suggest that they depend on partially distinct mechanisms [14,15].

In animal studies, insensitivity to negative outcomes has been mostly investigated using punishment procedures in which actions leading to a reward are contingently paired with electrical footshocks [9–13]. This punishment produces learning processes that result in the reduction of seeking and taking behaviors [15]. Importantly, differences in sensitivity to punishment have been used as a measure of transition from controlled to uncontrolled drug use and considered as a marker of addiction [11]. Obtaining a deeper knowledge of the behavioral and neurobiological basis of resistance to punishment is critical for better interpretation of results obtained using these procedures and for understanding the role of punishment processes in psychiatric disorders such as addiction [13,15].

The emotional impact of events that occur in the future is discounted (i.e. their perceived value decreases as a function of time) [31–33]. Differences exist among individuals in the rate of discounting, and individuals who discount reward more rapidly are considered impulsive and at risk to develop psychiatric disorders such as addiction [32]. Indeed, the negative consequences of compulsive and addictive behaviors often occur well after the execution of the compulsive action. Therefore, individuals that discount punishment too rapidly may not be able to appropriately consider the future consequences of their actions and may be particularly vulnerable to addictive disorders [34,35]. Importantly, punishments are temporally discounted in much the same way that rewards are less effective when delayed [33]. That is, people [36,37] and animals [38–41] are more willing to accept negative consequences when they occur after a delay. Therefore, as previously suggested [13,15], it is important to investigate the behavioral consequences of introducing a delay between a reward-seeking action and punishment.

We have recently developed a procedure to investigate punishment in rats that, similarly to what is commonly done in humans [16–18], individually calibrates the strength of punishment based on the animal’s behavior [19]. This self-adjusting progressive strength of shock (PSS) procedure allows obtaining PSS break points that quantify individual resistance to punishment [19]. Importantly, compared to other punishment procedures [20–29], the PSS procedure reduces exposure to high levels of shock that could be particularly aversive and persistently affect operant behavior [29], and it therefore could be considered a refinement in the 3Rs (replacement, reduction, refinement) principles of animal research [30]. Resistance to punishment in this procedure is sensitive to manipulations of motivation and appears to be both a trait (i.e. PSS break points are highly correlated under a wide variety of conditions) and a state (i.e. PSS break points are influenced by motivational states) [19]. Characterizing behavior in the PSS procedure may help in understanding mechanisms underlying resistance to punishment.

In this study, we used the progressive shock strength procedure to investigate the effects of a range of delays (0-12 sec) on the PSS break points. In brief, after training in a fixed-ratio 1 (FR 1) food procedure, rats were tested once per week for resistance to punishment in the PSS procedure or motivation for food in a progressive ratio (PR) procedure. Each delay was tested at least twice, and at the end of operant testing, we assessed anxiety-like behavior and pain sensitivity.

## Material and methods

Forty-eight male Sprague-Dawley rats aged 8-9 weeks (Janvier Labs, France), experimentally naive at the beginning of the study, were used in this study. The rats were divided into 2 cohorts of 24 rats which differed in the order of delay presentation. All experiments were conducted during the light phase and in accordance with European Union directives (2010/63/EU) for the care of laboratory animals and approved by the local ethics committees (COMETHEA).

### Food restriction

During operant procedures and until the end of the experiment, animals underwent food restriction to limit weight gain and to maintain operant behavior. Food (approximatively 20 g/day) was given 1 hour after the end of the experimental sessions, and rats had unlimited access to water for the entire duration of experiment.

### General experimental designs

Fig. S1 shows the experimental design for the two cohorts of rats. After 9 training sessions under fixed-ratio 1 (FR1) schedules, rats underwent PSS sessions with varying delays between animals’ lever presses and punishment delivery. For the first 3 PSS sessions, on alternate weeks, we also measured responding in the progressive ratio schedule. Afterwards, rats underwent only PSS test sessions, interspersed with normal FR1 schedules. The delay was fixed for a given session. Each delay was tested for 2 to 4 sessions. Tests were performed once per week, every fifth session (normally on Fridays).

Three different delays (3, 6 and 12 sec) were tested. The range of delays was chosen based on previous work showing that the ability of rats to learn the association between their behavior and a motivational stimulus depends on the temporal contingency of the response and the stimulus, and that beyond 8 sec this learning is impaired and aversion to footshock decreases to about 25% of initial value in a temporal discounting procedure [42].

In a first cohort of 24 animals, delays were presented in the following order: 0 sec (x 3 sessions), 3 sec (x2), 12 sec (x3), 6 sec (x3). To rule out an effect of the order of delays on the effects of shock, in the second cohort of 24 animals, delays were presented in a different order: 6 sec (x2), 3 sec (x2), 0 sec (x2), 12 sec (x2). 24-72h after the last operant sessions, we measured anxiety-related behaviors in an open-field and pain reactivity in the hotplate.

Since behavior did not differ significantly between the two cohorts, we combined the results.

### Food Reinforcement Apparatus and training procedure

Experimental chambers (MedAssociates, www.medassociates.com) were enclosed individually in sound-attenuation chests. Each experimental chamber had a recessed food tray, and two levers in the right wall. The floor consisted of bars that were connected to shockers (MedAssociates, ENV-414SA) that could deliver footshock, with electric current set to 0.45 mA. Each chamber was equipped with a food-pellet dispenser, which could deliver 45Lmg pellets to the food tray.

Experimental events were controlled by computers using MedAssociates interface and Med-PC IV software; Med-PC code used to conduct the procedures is available upon request. A diode light was present on each lever. One lever was assigned to be the active lever and the corresponding light was used as a conditioned stimulus for food reinforcement. A third diode light was installed on the opposite wall, and its flashing was used as a discriminative stimulus to indicate that food reinforcement would be associated with a foot-shock.

The general training schedule involved 45-min sessions of a schedule of food reinforcement in which one lever press (FR1) was required to obtain a 45-mg sucrose pellet. During these sessions, food availability was signaled by turning off the house-light, delivery of food was accompanied by flashing of the diode light above the lever for 2Lsec. Subsequently, the house light was turned on for an additional 18-sec time-out period, during which responding had no programmed consequences. Following the time out, a new trial started and the next response on the right lever was again reinforced. Responses on the inactive lever were recorded but never reinforced. Rats initially learned to respond for food during nine sessions under this schedule.

### Self-adjusting progressive punishment procedure

The self-adjusting progressive shock strength (PSS) procedure was the same as described by Desmercieres et al (2022). In brief, active lever presses resulted in the delivery of food rewards and foot-shocks of fixed intensity and variable duration. The self-adjusting procedure consisted of steps in which the shock duration was increased if the animal completed 2 trials in the previous step. The duration of the first step was 0 sec (no punishment), the second step was a low duration of 0.05 sec and subsequent shocks increased at each step for 20 steps. The durations of the steps were: 0, 0.05, 0.06, 0.07, 0.08, 0.09, 0.10, 0.12, 0.13, 0.15, 0.18, 0.20, 0.23, 0.27, 0.31, 0.35, 0.41, 0.47, 0.54, 0.62, 0.71 sec. If animals reached the final step, the duration of the shock was not further increased, and all subsequent shock were set at 0.71 sec. If rats did not emit any response for 5 min, shock duration was reset to 0 and the shock progression was reinitialized. The strength of the shock was measured by the electrical charge in millicoulombs (mC) that an animal was ready to receive to self-administer food pellets and was calculated by multiplying the fixed current of the shock (0.45 mA) by the duration in sec. The break point could be calculated by the total intensity of the shocks received during the session. We chose this parameter for the main analysis because it incorporates both the willingness to receive a given charge unit and the willingness to restart responding after an eventual punishment-induced pause.

### Progressive-ratio schedule

Under the progressive-ratio schedule of food reinforcement, the number of responses required to obtain a food pellet increased with each successive food pellet. The steps of the exponential progression were the same as those previously developed by Roberts and colleagues [43] adapted for food reinforcement [44,45], based on the equation: response ratio = (5e^(0.2^ ^×^ ^reinforcer^ ^number)^) − 5, rounded to the nearest integer. Thus, the values of the steps were 1, 2, 4, 6, 9, 12, 15, 20, 25, 32, 40, 50, 62, 77, 95, 118, 145, 178, 219, 268, 328, 402, 492, 603, and 737. Sessions under the progressive-ratio schedule lasted until 10 min passed without completing a step, which, under basal conditions, occurred within 1 h.

### Anxiety-related behaviors: Open Field

The open-field apparatus (Viewpoint, Lyon, France) consisted of a rectangular arena (50cm wide * 50cm long * 40cm high) of white plexiglass. After a 30-min habituation to the experiment room, rats were placed in the arena for 30 min. Their positions were recorded automatically by a camera and video tracking software (Viewpoint, Lyon, France). The software defined a virtual square (25cm * 25cm) delimiting the center zone and the border zone. Anxiety was measured as the percentage (%) of time spent in the center (time in the center/time in the border + time in the center * 100) so that more time spent in the center indicated a lower level of anxiety.

### Pain: Hot plate test

The hot-plate (Ugo Basile, model-DS 37) was maintained at 48 °C [46]. After a 10-min habituation to the experiment room, animals were placed into a glass cylinder with 25 cm diameter heated surface and 47 cm walls. The latency before escape or jumping was recorded. Experiments were stopped after a cut-off of 120s to prevent unnecessary pain or any tissue damage.

### Statistical Analysis

Med-PC data were analyzed using custom-made, freely available software written in Python, Med_to_csv (https://github.com/hedjour/med_to_csv) that uses raw data files to create complete tables for further analysis in GraphPad Prism. Data were checked for normality of distribution using the Shapiro–Wilk test. In our previous study, in the absence of a delay, PSS data did not show a normal distribution as a function of the electrical charge variable, but they did after natural logarithmic transformation of the variable. With the introduction of a delay, PSS data did not show normal distribution even after logarithmic transformation. Therefore, for statistical analysis, we used a nonparametric repeated measure Friedman rank test to analyze the effects of different delays in all rats.

To dichotomize individuals into shock sensitive or resistant groups we used the median split of the PSS break point measured by the total electrical charge sustained at delay 0 sec. Median splits were obtained in each cohort of rat separately. Whereas these choices are arbitrary, it should be noticed that similar results were obtained when using PSS break points at 6 or 12 sec of delay or when median splits were calculated merging the two cohorts.

For the investigation of the effects of delays in shock sensitive or resistant groups, we used two-way ANOVA for repeated measures with Geisser-Greenhouse correction to account for possible violation of sphericity, followed by Sidak’s post hoc test. Differences were considered significant when p < 0.05.

## RESULTS

### Food self-administration under basal conditions

Self-administration of food during the entire experiment is shown in Fig. S2 for the first cohort, and fig.S3 for the second cohort. Rats quickly learned to self-administer food under FR1 schedules. The initial baseline (BL), calculated as the mean ± SEM of responses per session during the three sessions before the first shock, was 121.74±1.53, and the final baseline calculated as the average of the last three sessions was 122.29 ± 2.45.

### Effects of delay on PSS break points

Increasing the delay from 0 to 3 did not produce significant increases in PSS break point. Only when the delay was further increased to 6 sec did the number of active responses significantly increase, from about 22 at delay 0 sec to about 48 at delay 6 and about 51 responses when the delay was set to 12 sec (Fig. 1A). The shock strength that rats were willing to receive increased from 0.97 to a maximum 8.17 mC at delay 12 Sec (Fig. 1B). This suggests that delay decreases the magnitude of the punishment effect.

**Fig. 1.**
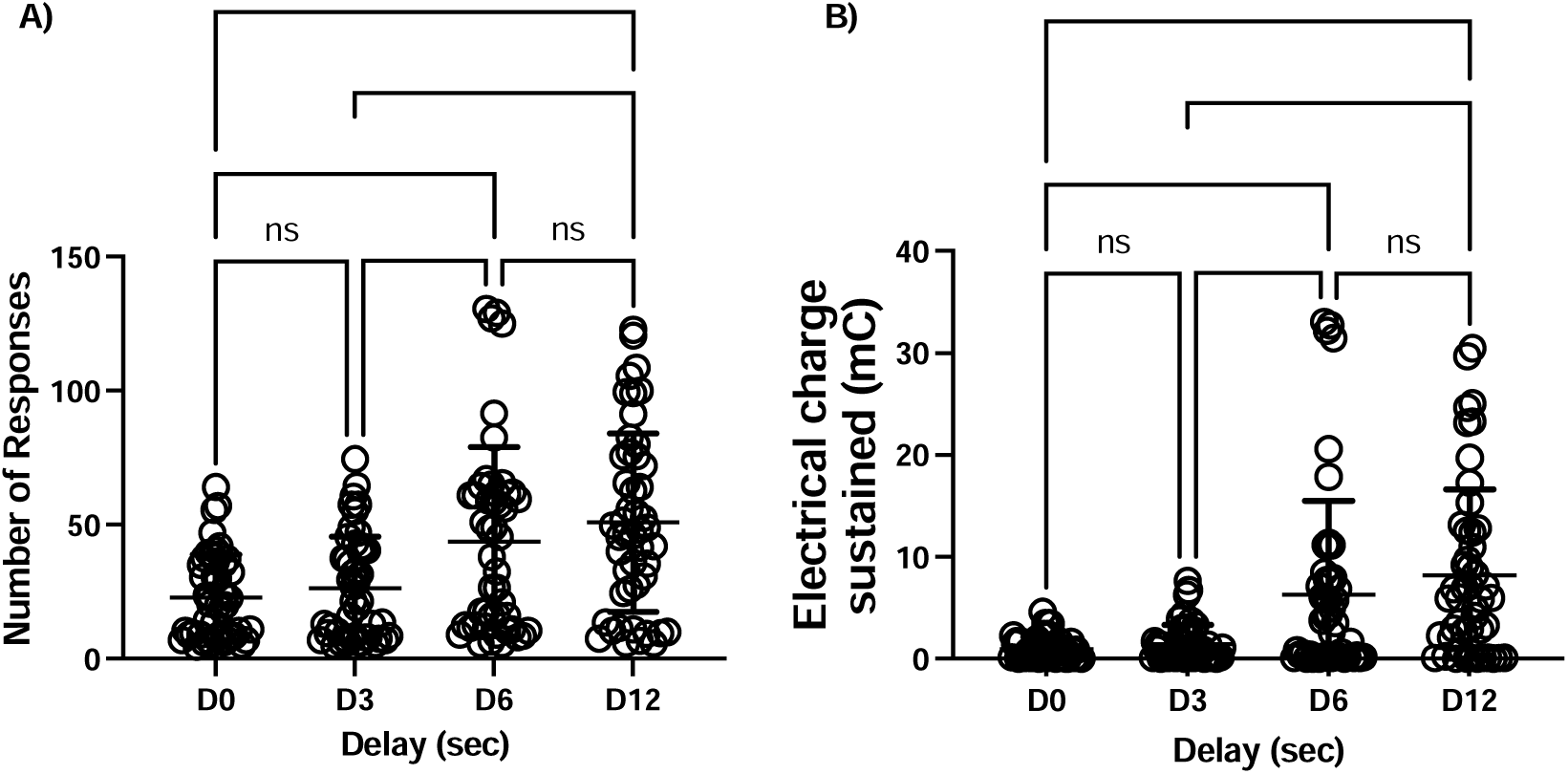
Effect of delay on resistance to punishment. Number of active responses (A) and PSS break point (total electrical charge sustained) (B) as a function of the delay between the response (and the food delivery) and the footshock. Data are expressed as mean ± SD of active responses (N = 48). Each data point corresponds to the average of at least 2 sessions at a given delay. Friedman nonparametric post- hoc test for repeated measures: ****, P < 0.0001.

### Correlation of PSS break points at different delays

One of our hypotheses was that individual differences in delay discounting of aversive consequences could interfere with PSS break points so that animals more resistant to immediate punishment would not necessarily be more resistant after introduction of a delay. Contrary to our prediction, correlation among PSS break points was very high and significant under all conditions (Fig. S4A; Spearman correlation r > 0.65 and p < 0.0001 for all values), suggesting that introducing a delay did not alter the relative sensitivity to punishment.

### PR break points and correlations with PSS break points

In the first weeks of the experiment, in parallel with PSS we also measured motivation in a PR procedure (Fig. S1). Consistent with our previous results [19], we found significant correlation among PR break points (Spearman correlation r > 0.32 and p < 0.05 for all values) but no correlation between PSS and PR breakpoints (Fig. S4A and S4B) confirming that these tasks do not measure exactly the same behavioral process [19,22,25,47].

### Identification of individuals with high or low resistance to shock-induced suppression

Visual analysis of PSS data in fig. 1 reveals that introducing a delay in the PSS procedures leads to an increase in the variability of behavior among subjects, with some animals showing increases in PSS with low 3 sec delays and others showing no or very little increases in the PSS break point even with 12 sec delays. Therefore, we decided to use a median split of PSS break points at delay 0 sec to classify animals as shock sensitive and shock resistant and to better characterize these subgroups. Day-to-day behavior during the entire duration of the procedure is shown in fig. 2 for cohort 1 and Fig. 3 for cohort 2. A few characteristics deserve to be highlighted. First, behavior in FR1 training session was very similar in the two groups during initial training but after repeated shock sessions, the number of responses in the sensitive groups decreased even on the first training session after the shock, with a slower return to baseline with repeated training sessions. This behavior suggests that shock-sensitive rats develop conditioned suppression of operant behavior. Second, responding of shock-sensitive and shock-resistant rats were clearly different in PSS sessions but were similar in PR sessions.

**Fig. 2.**
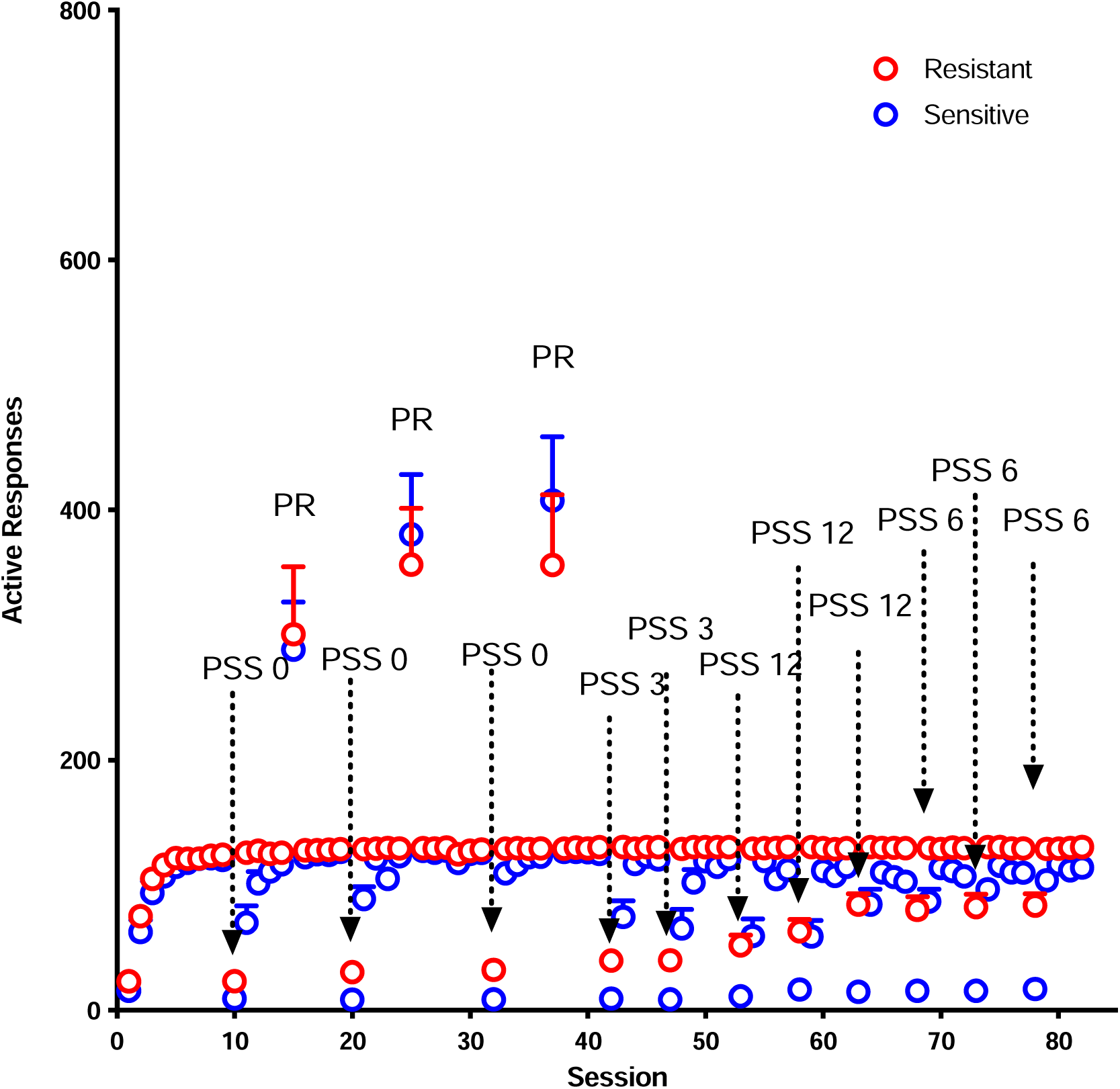
Operant behavior in shock sensitive and shock resistant rats during the entire experiment: cohort 1. Number of active responses in all the 84 sessions of the experiment for cohort 1. It should be noticed that during normal training and shock sessions each active responses produces the delivery of one sucrose pellet whereas in PR sessions each subsequent pellet requires an increasing number of responses. N = 12 per group.

**Fig. 3.**
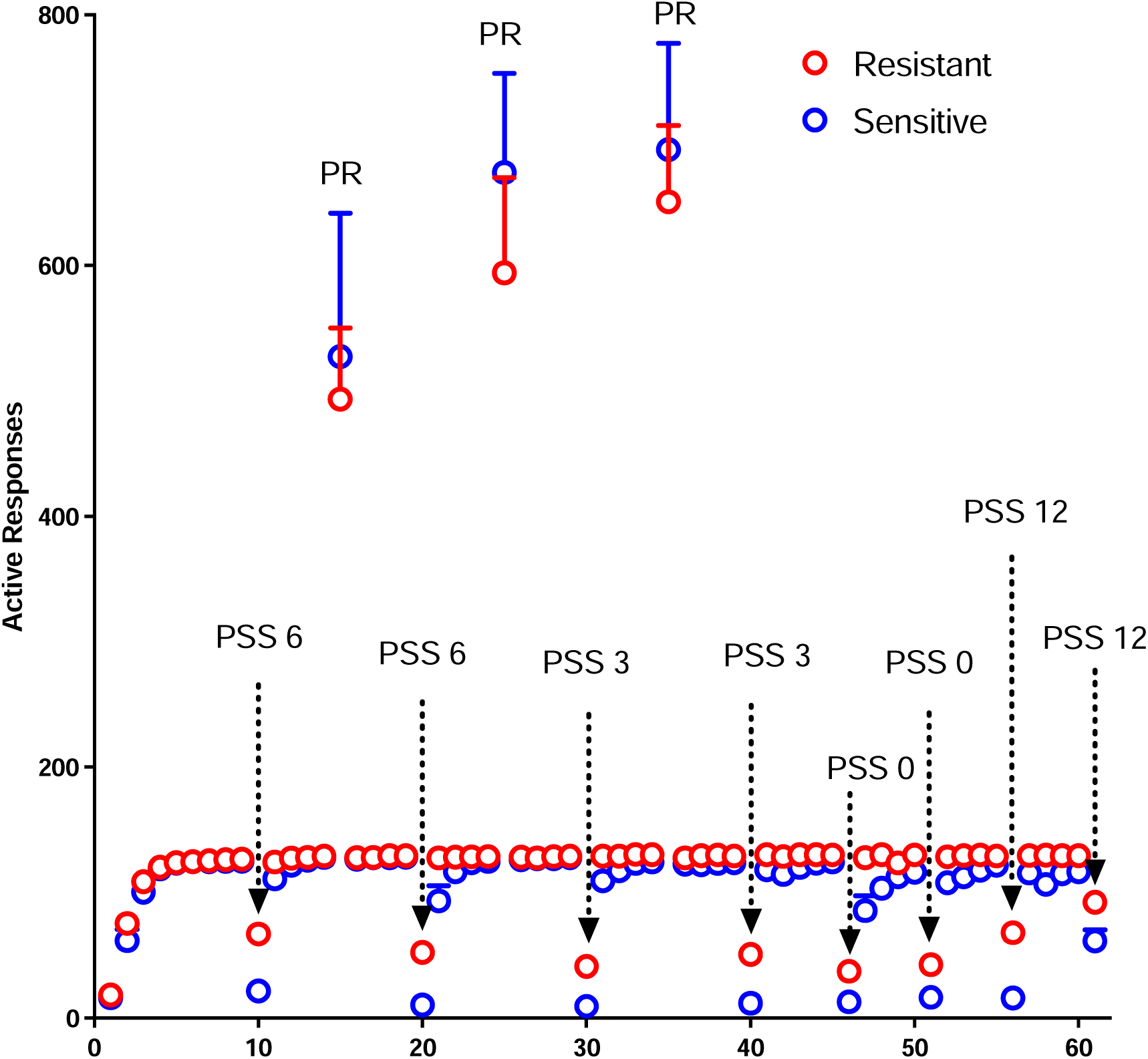
Operant behavior in shock sensitive and shock resistant rats during the entire experiment: cohort 2. Number of active responses in all the 61 sessions of the experiment for cohort 2. It should be noticed that during normal training and shock sessions each active responses produces the delivery of one sucrose pellet whereas in PR sessions each subsequent pellet requires an increasing number of responses. N = 12 per group.

### Effects of delay in high and low shock-resistant animals

PSS break point as a function of delay in sensitive and resistant animals is depicted in Fig.4. In both sensitive and resistant rats PSS break point increased with increasing delays; however, resistant rats showed significant increases with delays as low as 3 sec whereas sensitive rats showed increases only at 12 sec of delay. In addition, resistant rats showed significantly higher break points than sensitive rats at all delays. Statistical analysis revealed an effect of sensitivity to shock (Responses: F (1, 46) = 124.4 ; P < 0.0001; Electrical charge: F (1, 46) = 42.24 ; P < 0.0001), of delay (Responses: F (2.328, 107.1) = 43.10; P < 0.0001; Geisser-Greenhouse’s epsilon = 0.78; Electrical charge: F (1.999, 91.95) = 31.32; P < 0.0001; Geisser-Greenhouse’s epsilon = 0.67) and a sensitivity X delay interaction (Responses: F (3, 138) = 13.01, P < 0.0001; Electrical charge: F (3, 138) = 16.13; P < 0.0001).

**Fig. 4.**
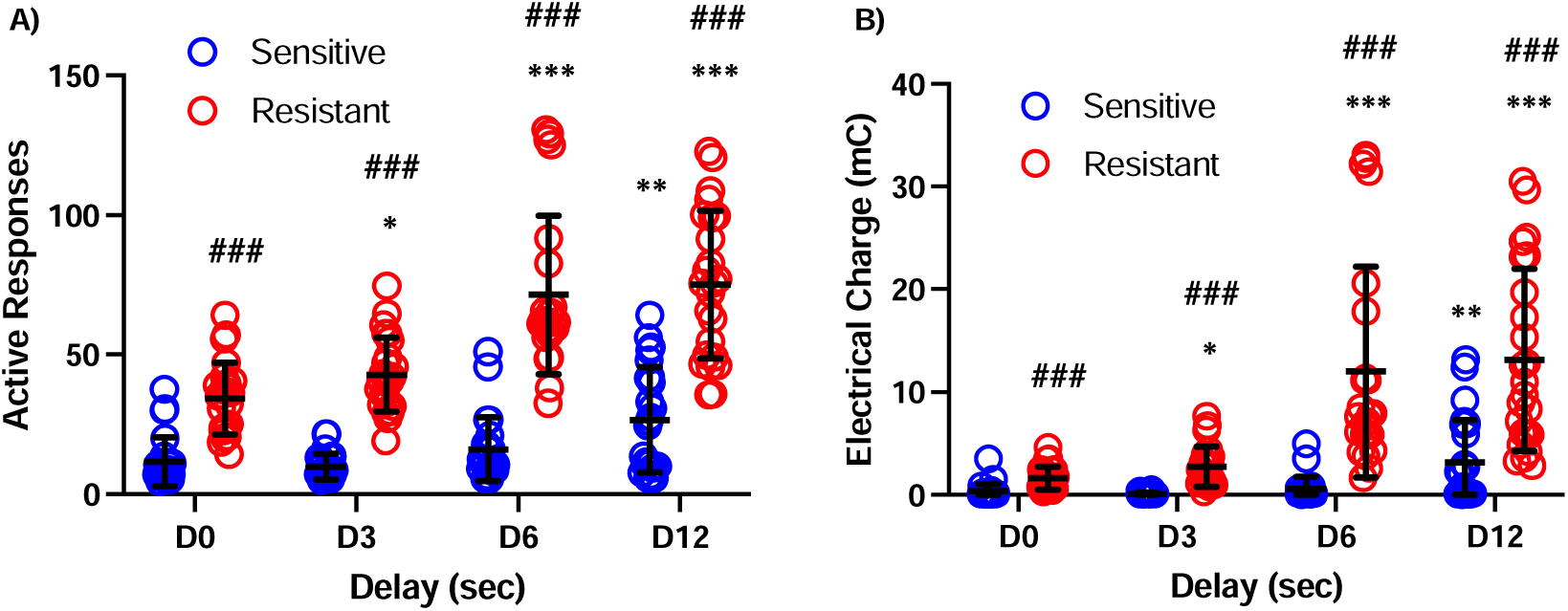
PSS break points as a function of the delay in shock-sensitive vs shock- resistant rats. PSS break point measured as the number of active responses (A) and total electrical charge sustained (B) as a function of the delay between the response (and the food delivery) and the footshock. Data are expressed as mean ± SD of active responses (Males n = 24 per group). Each data point corresponds to the average of at least 2 sessions at a given delay. Notice that median split was calculated separately in each cohort of rat which results in some overlapping of sensitive and resistant rats at delay 0. Two-way ANOVA for repeated measures: *, **, *** = P < 0.5, P < 0.001 and P < 0.0001 compared to Delay 0 sec (D0); ### = P < 0.0001 compared to shock sensitive rats.

### Conditioned suppression of operant behavior

As previously noted, after they started to receive shock training, shock-sensitive animals showed a reduction in the number of responses emitted even in training sessions in which shock was absent. Therefore, we calculated the difference between the day before and the day after the PSS session, and we called this measure the “suppression score”. Whereas resistant rats do not show any sign of suppression at any delay, sensitive rats show significant conditioned suppression that was similar at all delays (Fig. 5). Statistical analysis revealed a significant effect of sensitivity to shock (Responses: F (1, 46) = 44.92 ; P < 0.0001), no effect of delay and no significant sensitivity X delay interaction.

**Fig. 5.**
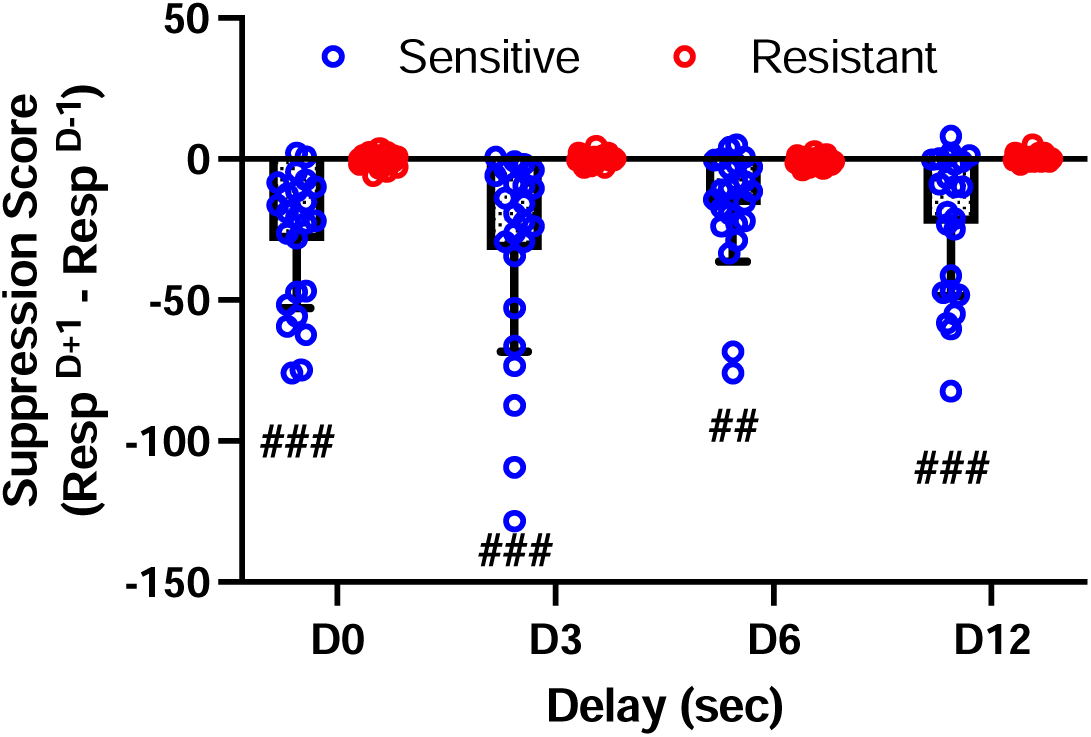
Conditioned suppression as a function of the delay in shock sensitive vs shock resistant rats. The suppression score was calculated as the difference in the number of active responses between the day after (D+1) and the day before (D- 1) the shock session. Data are expressed as mean ± SD of active responses (Males n = 24 per group). Two-way ANOVA for repeated measures: ##, ### = P < 0.001, P < 0.0001 compared to shock resistant rats.

### Anxiety and pain

Difference in sensitivity to shock could be due to trait-like differences in pain perception or anxiety. To determine whether these parameters influenced behavior, at the end of the operant sessions, we measured pain in a hot plate test and anxiety in an open field. We found that sensitivity to pain did not differ between sensitive and resistant rats (Fig. 6A). On the other hand, anxiety measured at the end of the experiments, was higher in sensitive compared to resistant rats (Fig. 6B, Mann-Whitney U = 164.5, P < 0.05).

**Fig. 6.**
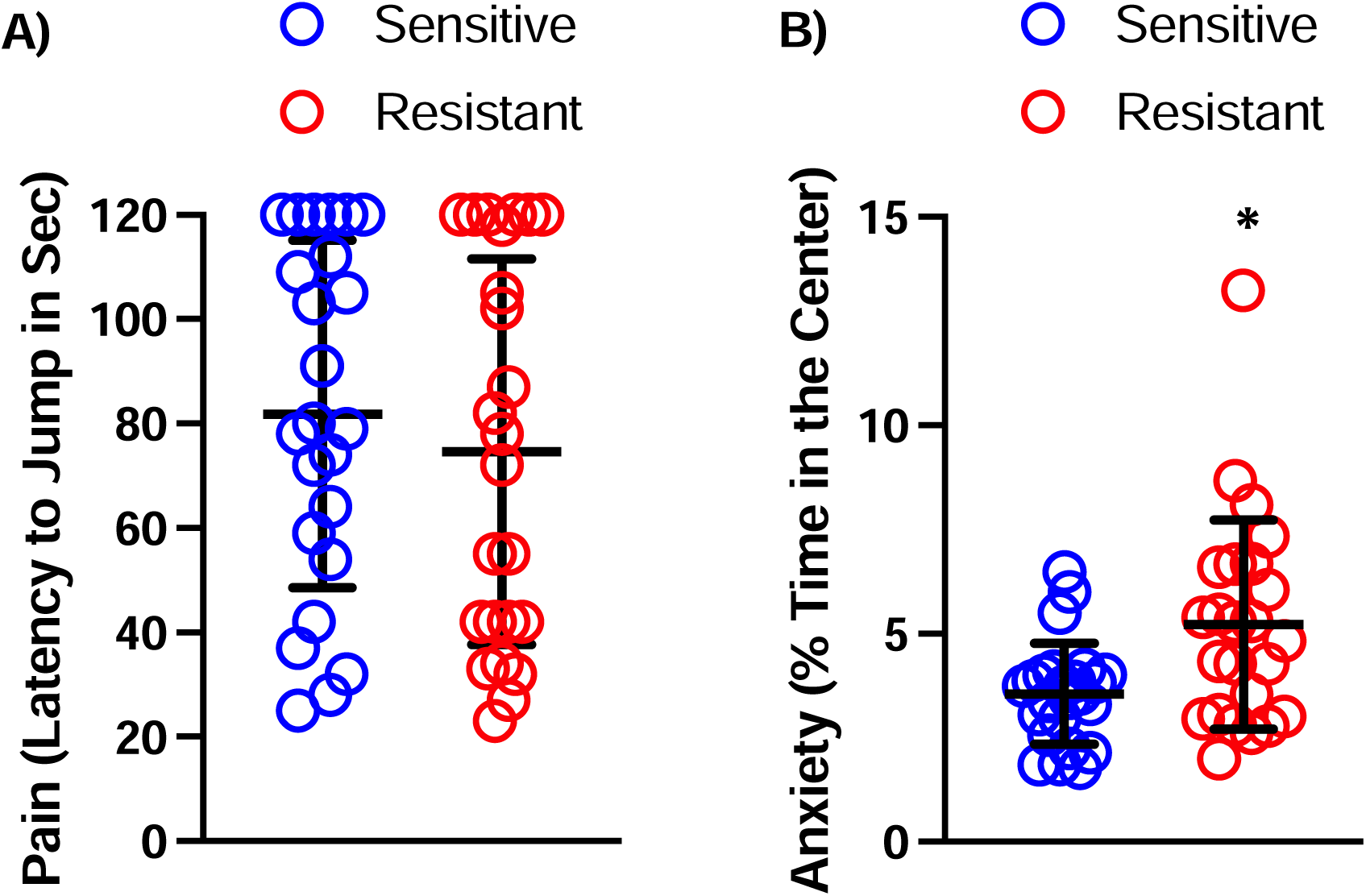
Pain sensitivity and anxiety in shock sensitive vs shock resistant rats. Pain sensitivity in the hot plate test (A) and anxiety-like behavior in an open field (B). These tests were performed at the end of the experiment. Note that the rats that spend more time in the center of the open field, are considered the less anxious. Data are expressed as mean ± SD of active responses (Males n = 24 per group). Mann-Whitney test: * = P < 0.05, P < 0.0001.

## Discussion

In this study, we investigated the effects of introducing a delay between reward and punishment in the PSS procedure [19]. When punishment was delayed, rats were willing to tolerate stronger shocks. Importantly, the introduction of a delay highlighted the existence of two populations of rats: one that not only is resistant to immediate punishment but that also discounts more future aversive consequences; the other that is sensitive to punishment and shows less temporal discounting. Finally, resistance to punishment was not associated with stronger appetitive motivation (as measured by the progressive ratio schedule) or with lower pain sensitivity, but it was negatively correlated with the development of conditioned suppression and anxiety-like behavior.

In humans, the negative consequences of maladaptive behaviors such as addiction are mostly delayed [34,35]. This delay contributes to the difficulty in adapting the behavior and reducing drug seeking and taking because, similar to what is commonly found with the positive value of rewards [48], the negative value of punishment diminishes with the delay [36–41]. In this study, consistent with those studies, we found that rats were willing to tolerate higher strengths of punishment when footshocks were delayed compared to those tolerated when punishment was immediate. However, consistent with previous work showing individual differences in delay discounting [41], we found considerable differences in the effects of delay on punishment. In fact, the rats that were more resistant to immediate punishment were also those that were more sensitive to temporal discounting. Therefore, introducing delaying punishment in the PSS procedures allows identifying a population that may be doubly vulnerable to developing compulsive-like behavior and addiction.

An unexpected finding in this paper is that shock-sensitive rats not only show a profound reduction in responding on shock days but also a significant reduction on the following days even in the absence of shock. This suppression disappeared and behavior tended to return to baseline only after several sessions without shocks. Thus, the emotional impact of footshock appeared to be a major contributor to the effects of punishment in this study. Using a procedure that separates pavlovian conditioning from operant punishment, Jean-Richard-Dit-Bressel and colleagues showed that under certain conditions, conditioned fear and punishment can be dissociated [49]. In contrast, in the PSS procedure, pavlovian and operant conditioning are not separated, which causes a strong association between punishment and conditioned suppression. Future studies will be needed to investigate whether behavior in the PSS procedure and the procedure of Jean-Richard-Dit-Bressel overlap and whether these procedures identify the same populations of resistant and sensitive rats.

In humans, some individuals could use drugs while managing to maintain control and whereas others lose control over the drug and become addicted [50,51]. However, it is difficult to predict who will develop addiction and who won’t. Animal studies can provide useful information about the factors that may increase the risks to develop compulsive-like behavior. For drugs, after 20-30 sessions but not earlier, compulsive-like behavior develops in a minority of rats [21,52]. Conversely, in mice models of food addiction, resistance to punishment using conventional fixed-intensity procedure did not increase and it even decreased over time [53,54]. In this study, we tested PSS procedure over long periods (4-6 months) and in a high number of sessions (>80 sessions for cohort 1 and >60 sessions for cohort 2) and we found no evidence of a progressive loss of control over food seeking. Indeed, animals identified as shock-sensitive or shock-resistant after a few sessions, continued to show similar behavior for the entire duration of the experiment. This suggests that at least for food reward, sensitivity to punishment in the PSS procedure is an individual trait that is not appreciably affected by repeated exposure to food reward.

Previous studies have shown that experience with punishment can profoundly affect behavior in future situations [29,55]. In particular, early experience with a high level of shock may induce long-lasting hypersensitivity to shock and suppress behavior in a persistent manner [29]. Conversely, experience with low levels of shock can induce tolerance to punishment and allow individuals to tolerate higher levels of shocks later on [55]. The PSS procedure follows in the latter category. In the present study, changes in PSS were confounded by introduction of delays, but unpublished results in our lab show that upon repeated exposure, rats reach slightly higher PSS break points compared to initial levels. However, we found that PSS break-points are highly correlated throughout the experiments, suggesting that whereas absolute individual levels may increase, relative resistance to punishment in the PSS procedure is relatively stable over time, regardless of the order of delays to which animals were exposed. This is a potentially useful feature of the PSS procedure.

Anxiety and addiction are often comorbid psychiatric disorders and their relationships are bidirectional [56]. In this study, we found that anxiety measured in the open field was higher in shock -sensitive than in shock-resistant rats. If we consider shock -resistant rats as addiction-prone, this result is apparently at odds with previous findings in humans and animals. However, it should be noticed that we measured anxiety at the end of the experiment and therefore, the results can be affected by previous training in the PSS and could therefore be the consequence rather than the cause of different behavior in the PSS. Indeed, anxiety in sensitive rats was associated not only with lower PSS break points but also with higher conditioned suppression, suggesting that repeated experience of fear in an operant context may have led to diffuse fearful behavior. Importantly, in a previous study, PSS break point and anxiety were not correlated [19]. The main difference between that study and this one is the introduction of delay suggesting that temporal degradation of contingency induced an anxious phenotype. These results are reminiscent of previous studies in which probability degradation of punishment contingency produced anxiety-like behavior [57,58]. Importantly, in our study shock-sensitive rats showed higher anxiety than shock-resistant rats, suggesting that these animals became afraid of their environment even when no real threat was present.

Depending on the circumstances, being relatively resistant or relatively sensitive to punishment could be either adaptive or maladaptive. In our procedure, we found a subset of shock-resistant rats that kept seeking food, but they still adapted their behavior depending on the intensity of the shock or the delay. In contrast, the subset of shock-sensitive rats always stopped responding for food after a few shocks, and their behavior changed little when the punishment was delayed. In addition, sensitive animals showed conditioned suppression in the absence of punishment, a behavior that can hardly be considered optimal in the case of food reward. More generally, shock-sensitive and shock-resistant animals appear to have different strategies to cope with environmental challenges. At the individual level, the conservative (shock-sensitive) strategy may be more adaptive if risks are maintained and frequent whereas the risk (shock-resistant) strategy may be more adaptive if risks are temporary and rare.

In conclusion, introducing a delay between reward and punishment, we identified a subpopulation of rats that was highly sensitive to shock regardless of delay, to the extent that their responding was suppressed (and food reward was lost) even when shock was discontinued. These same individuals also showed anxiety-like behavior in a novel environment and may represent a population vulnerable to opportunity loss. In contrast, the rest of the rats showed resistance to punishment and this phenotype was even more pronounced when the consequences of food seeking were delayed. These individuals represent a population that would be particularly at risk to develop addiction-like behavior. Thus, the PSS procedure identifies factors that influence resistance to punishment within a behavioral economics paradigm, and future work should determine whether it can be used to assess vulnerability to develop drug addiction.

## Acknowledgements

This work was supported by the Centre National pour la Recherche Scientifique, the Institut National de la Santé et de la Recherche Médicale, the University of Poitiers, the Nouvelle Aquitaine CPER 2015-2020 / FEDER 2014-2020 program “Habisan” and the Nouvelle Aquitaine grant AAPR2020A-2019-8357510 (PI : M. Solinas) and the IRESP/Inca grants « IRESP-19-ADDICTIONS-20 » (PI : M. Solinas) and « IRESP-AAPSPA2021-V1-07 » (PI : N. Thiriet). The contribution of LVP was supported by the Intramural Research Program of the NIH, National Institute on Drug Abuse.

We thank Pauline Belujon for helpful comments on a previous version of the manuscript.

This study has benefited from the facilities and expertise of PREBIOS platform (Université de Poitiers).

