## Supplementary material for "Delay of punishment highlights differential vulnerability to developing addiction-like behavior toward sweet food": Fig. S1

### Slide 1
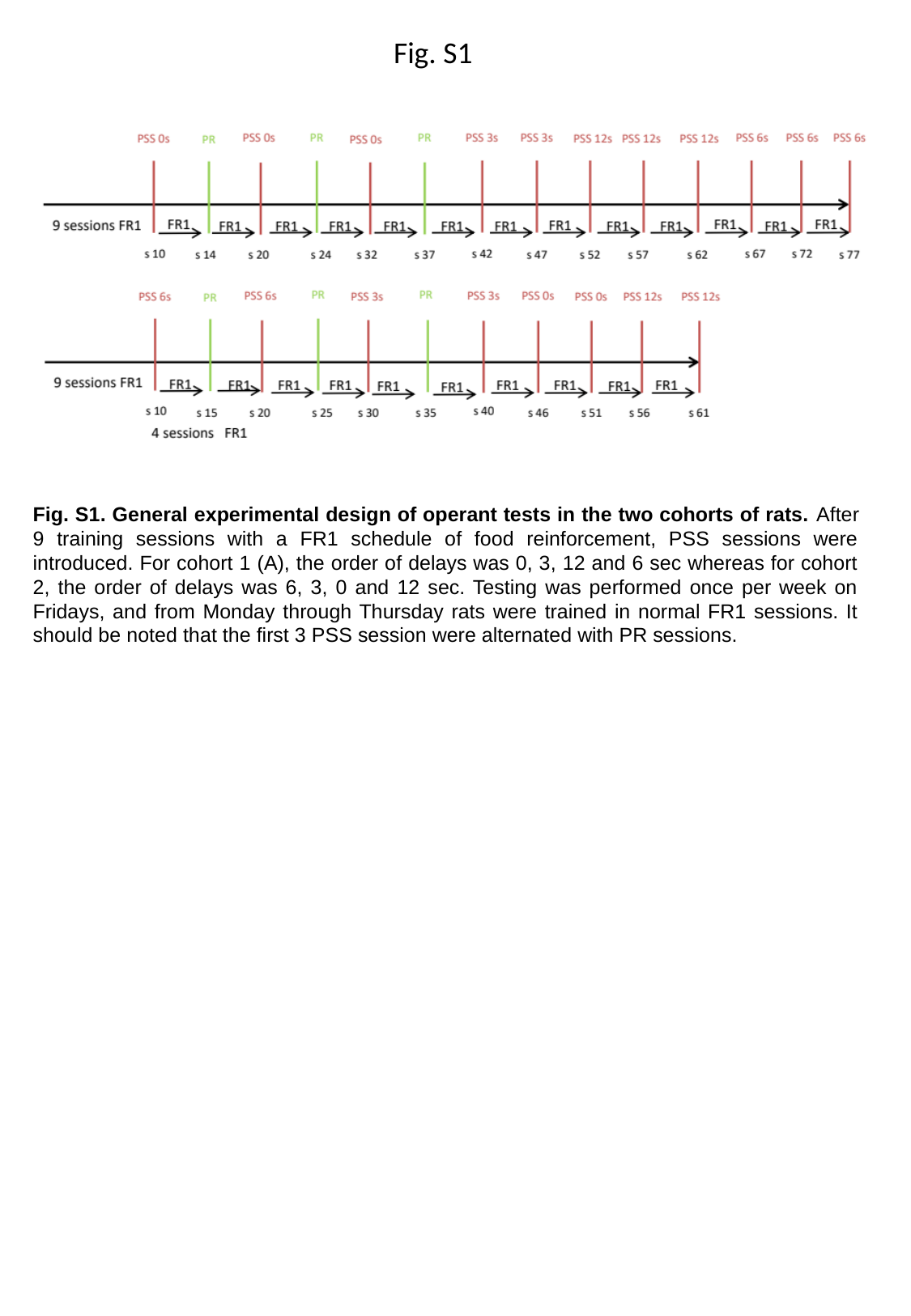

Fig. S1
Fig. S1. General experimental design of operant tests in the two cohorts of rats. After 9 training sessions with a FR1 schedule of food reinforcement, PSS sessions were introduced. For cohort 1 (A), the order of delays was 0, 3, 12 and 6 sec whereas for cohort 2, the order of delays was 6, 3, 0 and 12 sec. Testing was performed once per week on Fridays, and from Monday through Thursday rats were trained in normal FR1 sessions. It should be noted that the first 3 PSS session were alternated with PR sessions.

### Slide 2
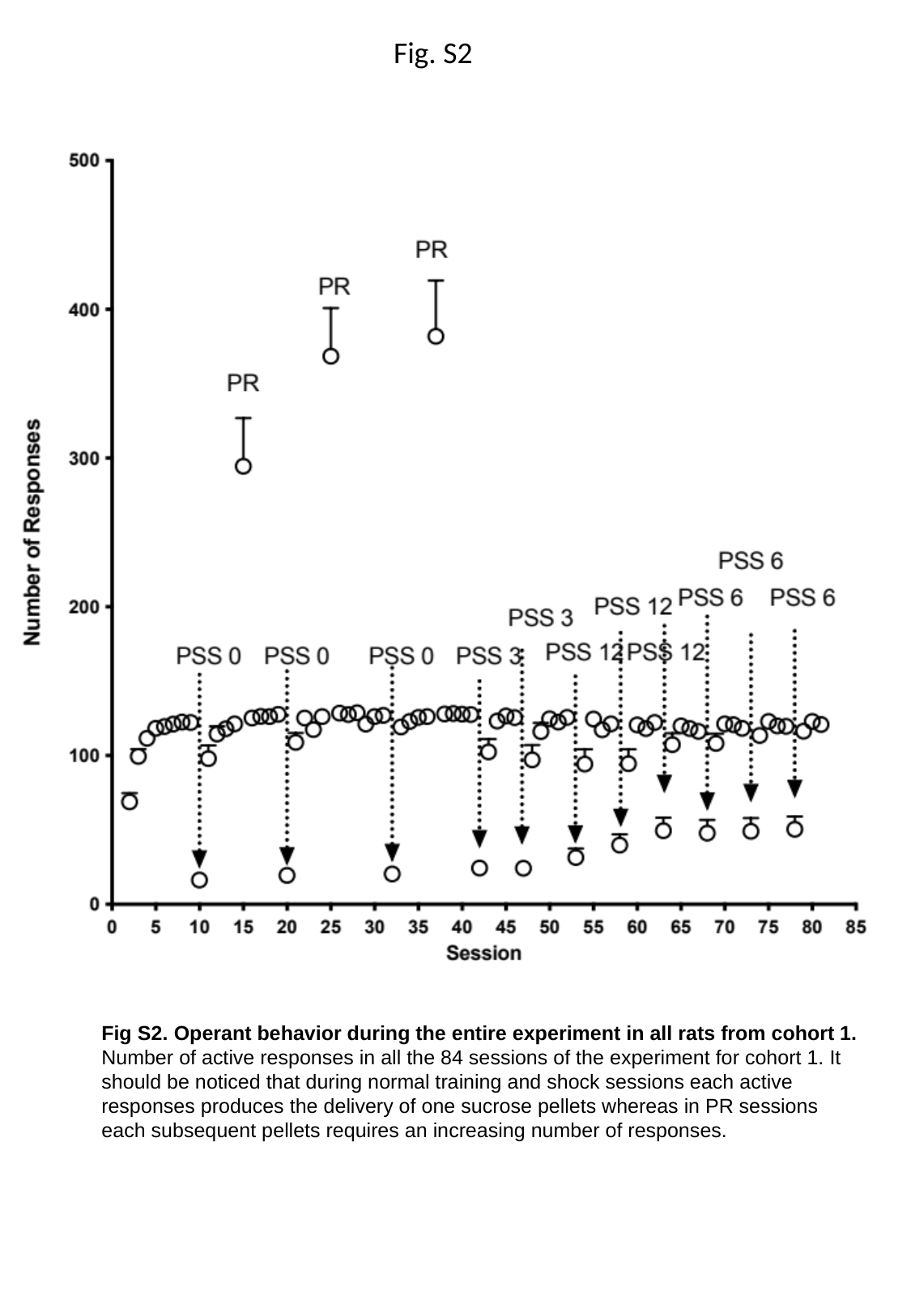

Fig. S2
Fig S2. Operant behavior during the entire experiment in all rats from cohort 1. Number of active responses in all the 84 sessions of the experiment for cohort 1. It should be noticed that during normal training and shock sessions each active responses produces the delivery of one sucrose pellets whereas in PR sessions each subsequent pellets requires an increasing number of responses.

### Slide 3
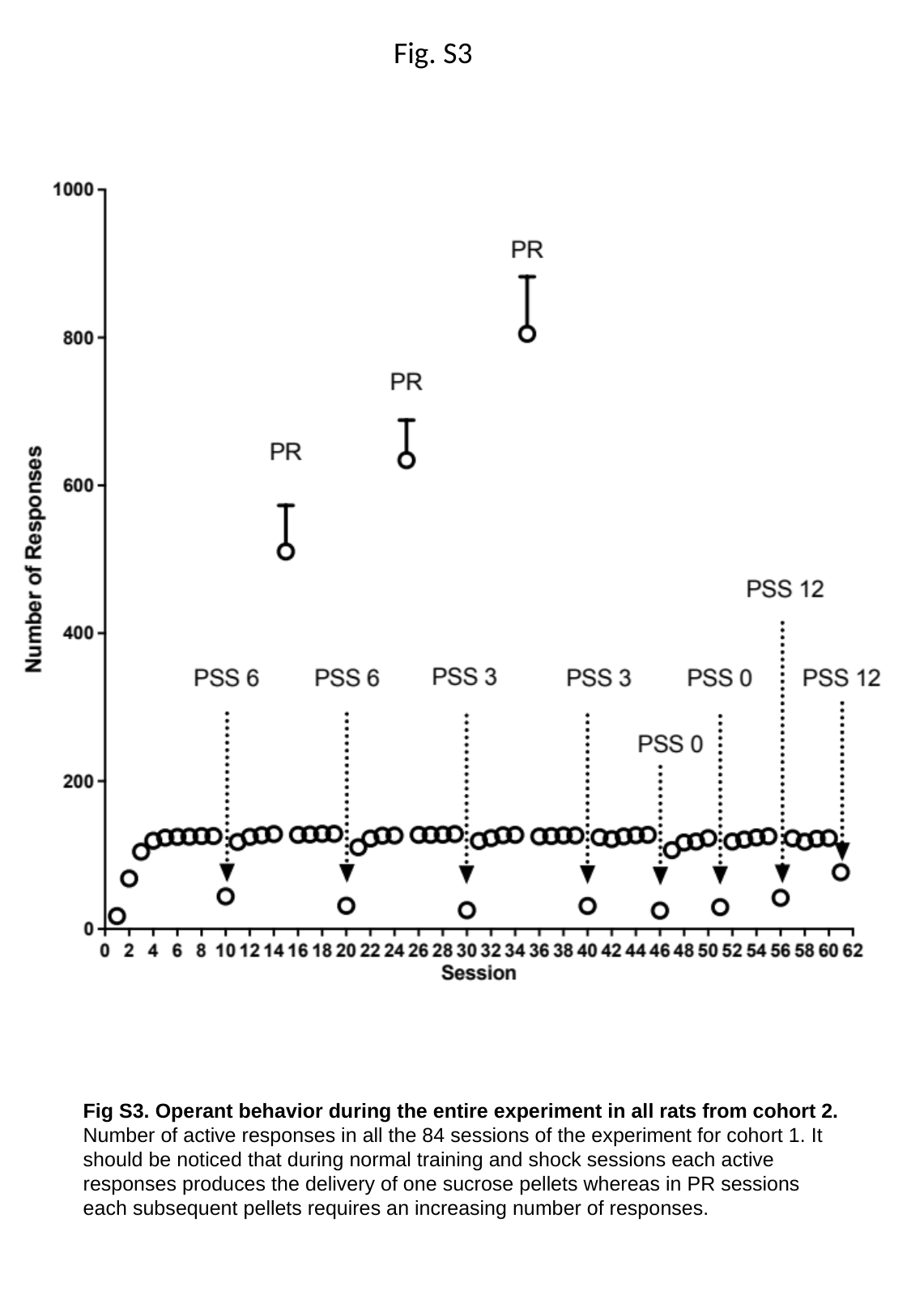

Fig. S3
Fig S3. Operant behavior during the entire experiment in all rats from cohort 2. Number of active responses in all the 84 sessions of the experiment for cohort 1. It should be noticed that during normal training and shock sessions each active responses produces the delivery of one sucrose pellets whereas in PR sessions each subsequent pellets requires an increasing number of responses.

### Slide 4
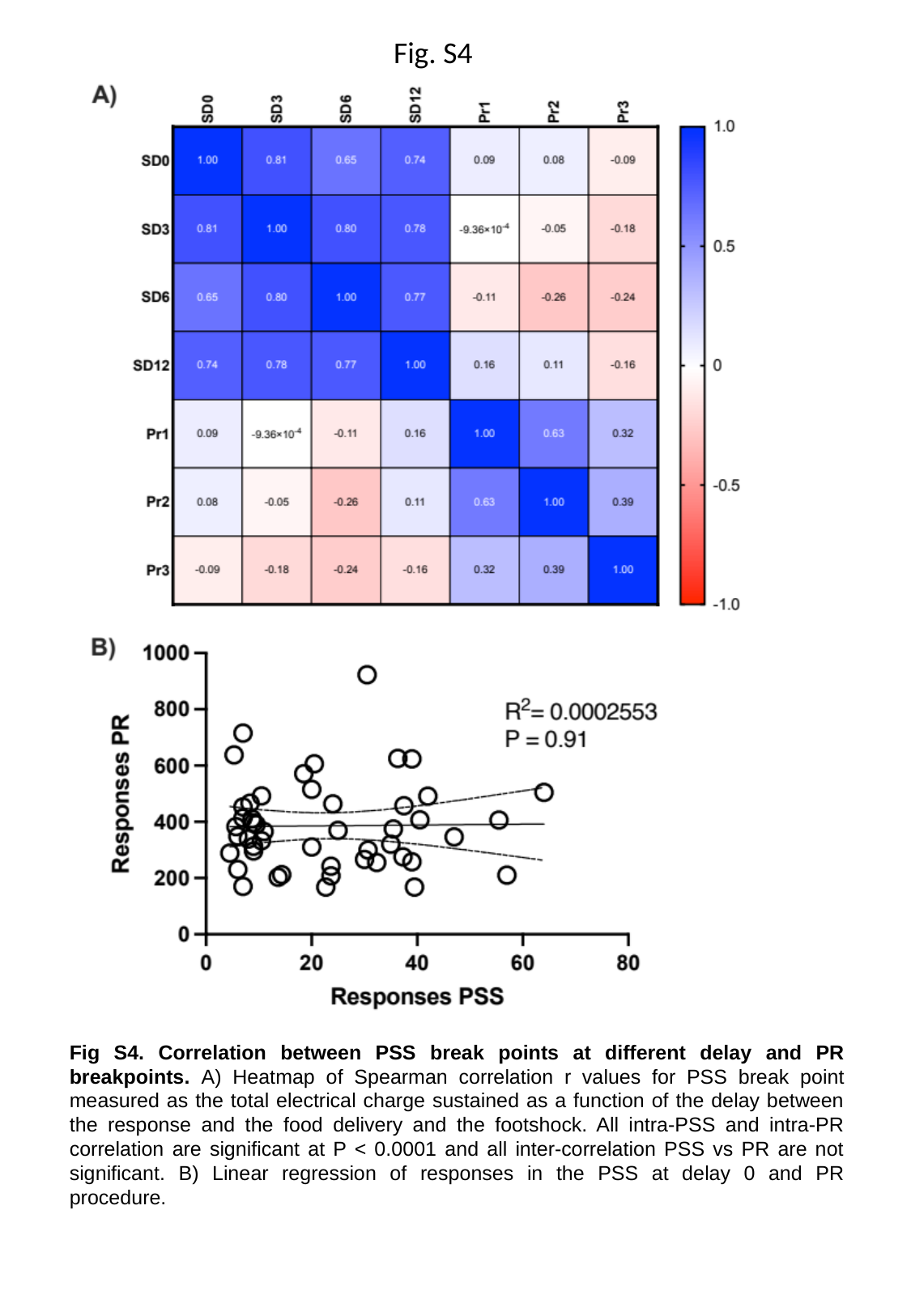

Fig. S4
Fig S4. Correlation between PSS break points at different delay and PR breakpoints. A) Heatmap of Spearman correlation r values for PSS break point measured as the total electrical charge sustained as a function of the delay between the response and the food delivery and the footshock. All intra-PSS and intra-PR correlation are significant at P < 0.0001 and all inter-correlation PSS vs PR are not significant. B) Linear regression of responses in the PSS at delay 0 and PR procedure.
